# Exploring the latent space of transcriptomic data with topic modeling

**DOI:** 10.1101/2024.10.31.621233

**Authors:** Filippo Valle, Michele Caselle, Matteo Osella

## Abstract

The availability of high-dimensional transcriptomic datasets is increasing at a tremendous pace, together with the need for suitable computational tools. Clustering and dimensionality reduction methods are popular go-to methods to identify basic structures in these datasets. At the same time, different topic modeling techniques have been developed to organize the deluge of available data of natural language using their latent topical structure.

This paper leverages the statistical analogies between text and transcriptomic datasets to compare different topic modeling methods when applied to gene expression data. Specifically, we test their accuracy in the specific task of discovering and reconstructing the tissue structure of the human transcriptome and distinguishing healthy from cancerous tissues. We examine the properties of the latent space recovered by different methods, highlight their differences, and the pros and cons of the methods across different tasks. Finally, we show that the latent topic space can be a useful embedding space, where a basic neural network classifier can annotate transcriptomic profiles with high accuracy.

## 1 Introduction

The number of large-scale and high-dimensional datasets of gene expression obtained with RNA sequencing is growing at a frantic pace [49, 15]. This data accumulation generates a parallel growing need for computational methods to identify the hidden relevant structures [8, 31]. For example, the efficient identification of patterns in expression data is a key step in the development of a precision medicine program, ranging from the discovery of disease-specific expression profiles for diagnosis to the stratification of patients [2]. Analogously, in the context of single-cell RNA sequencing data, one often needs to identify the different cell types and their expression hallmarks, and this hierarchical structure of cell types has to be inferred in the presence of several confounding factors [21].

For all these tasks, the number of different algorithms and tools that have been developed and applied to RNA sequencing data is huge [22]. For example, clustering algorithms are classic go-to methods to find groups at the level of samples (e.g., patients or single-cells). At the gene level, clustering can instead be used to extract the genes whose expression is collectively orchestrated in a particular system or in response to treatment.

Dimensionality reduction is another key step in many standard computational pipelines. For example, Principal Component Analysis or analogous non-linear methods, such as t-SNE, are widely used for both data-preprocessing and visualization [19, 20, 2]. The underlying hypothesis is that the number of relevant variables needed to efficiently describe the system is well below the number of genes, and thus concise but informative representations can be built [4]

This paper analyzes a class of algorithms, called topic models, that can essentially provide fuzzy clustering and dimensionality reduction simultaneously in a probabilistic framework [5]. Topic models were developed to organize texts of natural language using a space of latent variables, i.e. the topics, that define the different word usage probabilities, and can be interpreted as a more compact, low dimensional description of the dataset. Topics can be used to define a graded membership structure, or a fuzzy clustering, of texts in a corpus. While originally developed for natural language processing, topic modeling is recently finding applications in biology, and more specifically in genomics [64, 61, 54, 8, 48, 55, 56, 27, 18, 41]. Indeed, genomic data share several fundamental statistical laws with natural language and other modular complex systems [29, 25, 28].

However, the topic structure can be inferred using different methods that are based on alternative assumptions and priors. It is not clear a priori what are the pros and cons of each method, in particular in the genomic context. This is our focus in this paper. We will examine different approaches on simple and clear-cut benchmark datasets from RNA sequencing experiments. We will compare the performances of different algorithms in terms of clustering of samples and genes and analyze how their specific statistical priors affect the results in the context of genomic data. We will also discuss in detail the interpretability and biological relevance of the latent variables extracted by different topic modeling techniques.

The most popular topic modeling method is the Latent Dirichlet Allocation (LDA) [5]. It assumes that documents are generated from a mixture of topics, and each topic is characterized by a distribution over words. Iterative algorithms are used to infer the underlying topics and their distributions within the corpus. LDA employs Dirichlet priors to model the distribution of topics within documents and the distribution of words within topics. Such priors force few words to be representative of a topic, and analogously few topics to characterize each document. This choice is mostly motivated by mathematical and computational convenience, but it is not clear how appropriate it is in the genomic context, and what are its consequences. LDA has been previously applied to RNA sequencing data for data exploration [8], successfully retrieving genes whose expression can distinguish samples from different tissues.

Topic modeling has been recently connected to the problem of community detection in networks [10], and a new algorithm based on the hierarchical Stochastic Block Model (hSBM) has been formalized with the ambition to solve some of the weaknesses of LDA [12]. We have recently shown how this method can be used in the context of cancer genomics [55, 56], but a clear comparison with other topic modeling techniques in different genomic contexts is still needed.

Besides hSBM and LDA, we will also consider an alternative topic modeling approach called Topic Mapping (TM) [23], which has not been tested so far in the genomic context. This method uses the co-occurrence of words to create a network of tokens. Topics are then defined as basic network structures.

Topic modeling techniques will be also compared to more classic and popular computational methods such as the Weighted Gene Correlation Network Analysis [24] (WGCNA). WCGNA is not formally a topic modeling method but still has several commonalities with this class of methods. In fact, WGCNA is widely used to search for correlated groups of genes, which are called gene “modules”, that can be loosely associated to topics. WGCNA does not provide a natural way to define clusters or groups of samples, but standard clustering techniques can be applied using the modules (i.e., the topics) as features.

Finally, we will also consider standard clustering methods such as the well-known hierarchical clustering algorithm [17]. With hierarchical clustering, we can search for groups of samples based on their relative Euclidean distance in the feature space. We can thus compare these groups to the clusters that different topic modeling methods identify. This comparison will provide a better understanding of the advantages of complex topic inference with respect to easy-to-interpret classic methods.

We will show that different algorithms generally find different topic distributions (or embedding spaces) with alternative structures and statistical properties. In general, there is not an optimal algorithm for every task [47]. Our goal is to highlight the potential biases and tendencies that affect different algorithms. In fact, even in presence of comparable performances on a specific task, different approaches can identify alternative structures and show different biases [43]. This finding represents an issue that has to be considered in the context of interpretability and causal inference. Interestingly, the topics we find with alternative methods, although often poorly overlapping, are generally well correlated with the biological properties of the samples analyzed.

Topic models also provide a low-dimensional space in which the samples can be represented. A simple neural network can be trained on this low-dimensional topic space with the task of labelling new samples. Specifically, a classifier in this space can distinguish samples from different tissues, as well as detect diseased samples over healthy ones with very high accuracy despite the dimensionality reduction.

## 2 Results

### 2.1 Discovering the structure of the transcriptome of human tissues with different topic modeling approaches

We start the analysis by applying different topic modeling techniques to a large-scale dataset of RNA-sequencing samples of healthy human tissues from the Genome Tissue Expression project (GTEx) [30]. The performance of topic models and commonly used clustering methods can be compared to the simple task of tissue separation. Hoverwer, topic models have a probabilistic nature that provides an additional layer of information.

Figure 1 reports the results obtained using the topic modeling technique called hSBM [12]. The output of this model consists of blocks of samples (which we will denote as “clusters”) on one side, and blocks of genes (i.e., the topics) on the other side of a bipartite network. As a probabilistic method, these blocks can be fuzzy and eventually overlap. However, at this stage, we focus on not overlapping blocks to allow a straightforward comparison with clustering methods. Topic models also provide a probabilistic membership of samples to the topics. A mixture of topics can describe a sample via the probability distribution *P* (topic|sample) defined in Eq. 3 of the Methods section. Analogously, the gene membership to topics is probabilistic. The distribution *P* (gene|topic), defined in Eq. 4, captures the relative contribution of a gene to a specific topic. Moreover, the hSBM model has a hierarchical structure. The blocks can be organized in *L* layers, and blocks of layer *l* are nodes in layer *l* + 1 [35].

**Figure 1:**
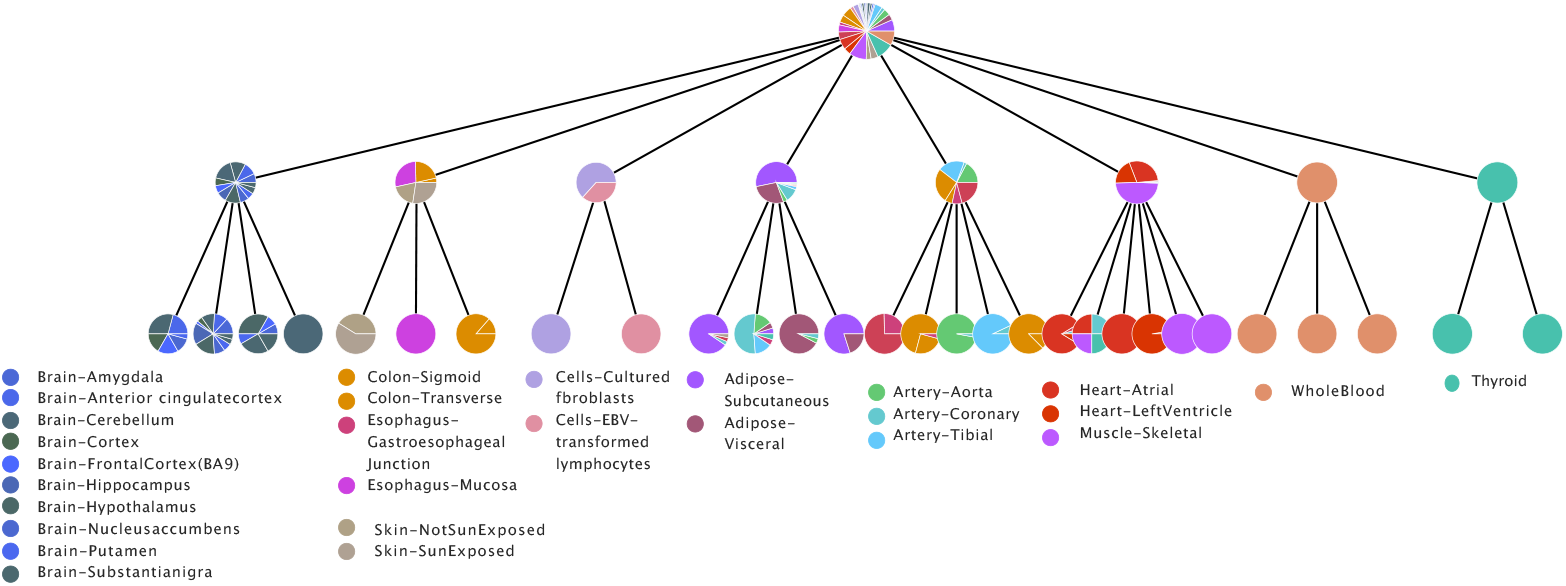
The hierarchy structure is represented as a tree. The first layer from the top is an all-samples node by construction. Then the algorithm found another node with all samples, at this point only the gene side of the network is partitioned. Then the algorithm found 8 clusters with a rough separation of tissues. From left to right the reader can identify a Brain cluster, a cluster with Colon, Esophagus, and Skin, a cluster with the label Cells (i.e., cultured fibroblasts and EBV-transformed lymphocytes), one mostly with Adipose tissue, one with Artery, one with Muscle and Hearth, one of Blood, and in the end one with Thyroid. In the next layers, the classification becomes finer, from Brain a Brain - Cerebellum cluster emerges, Skin and Colon are separated, Cells are split into Fibroblasts and Lymphocytes, Aorta, Coronary, and Tibial emerge, Hearth and Muscle are separated.

In principle, topic modeling algorithms should first recognize the basic tissue structure hidden in the expression profiles, and then recognize more fine-grained structures such as the sub-tissues. Figure 1 suggests that indeed this is roughly the case.

Besides hSBM, we ran the same analysis with other algorithms, specifically the well-known model LDA [5] and the more recent Topic Mapping (TM) [23]. We also compared the cluster structure found by WGCNA [24, 63], which outputs *modules* of correlated genes, by using the standard implementation in its R package https://cran.r-project.org/package=WGCNA. The Methods section reports the implementation details for each algorithm. Finally, we also tested hierarchical clustering on the same task [17]. We included it in the comparison since it is widely used thanks to its simplicity.

To reduce the complexity of the problem, we selected a subset of GTEx with 1 000 samples from the 10 most represented tissues, and 3 000 highly variable genes, as explained in more detail in the Methods section. In this setting, the output of hSBM has four layers with 1, 8, 29, 978 clusters at the sample level, and 22, 273, 2 725 and 2 880 topics, on the other side of the network. The first and last partitions are clearly not informative, while the two intermediate ones (8 and 29 clusters) are expected to capture the actual organization in tissues of the samples. In particular, the first one has a number of clusters close to the actual number of tissues in input, while the second partition (29 clusters) is expected to capture the subdivision of each tissue in more fine structures, for example distinguishing the cortex from the cerebellum in brain samples. Notice that the first partition of the genes into 22 topics happens when the samples are still all in the same cluster. This could be due to the asymmetry of the network. In fact, the number of genes is three times the number of samples. The second partition at the gene level, defining 273 topics, corresponds instead to the partition of samples in tissues.

hSBM is fully non-parametric: the hierarchical structure defined by the number of clusters and layers is automatically selected by the model [35]. Interstingly, the inferred hierarchical organization agrees with the hierarchical organization of tissues. For instance, from the cluster composed by *Brain* samples the following splitting leads to a sub-cluster composed by the *Brain - Cerebellum* samples. The possibility of separating brain samples from expression information was indeed previously observed by [30], when the dataset was released. Other examples of tissue and subtissue separations are reported in the caption of Figure 1.

We compare the performances of the different algorithms in Figure 2 using the Normalised Mutual Information (*NMI*) as a measure of the clustering quality. This measure was proposed by [50] to evaluate topic modeling performances on synthetic corpora. Since the *NMI* is not null for a random partition (there may be some residual entropy), we choose to scale the *NMI* score to a simple random null model obtained by reshuffling labels and fixing the number and the size of clusters. We will refer to this normalized measure as *NMI* ^*^. The Methods Section reports all the details about the evaluation metrics.

**Figure 2:**
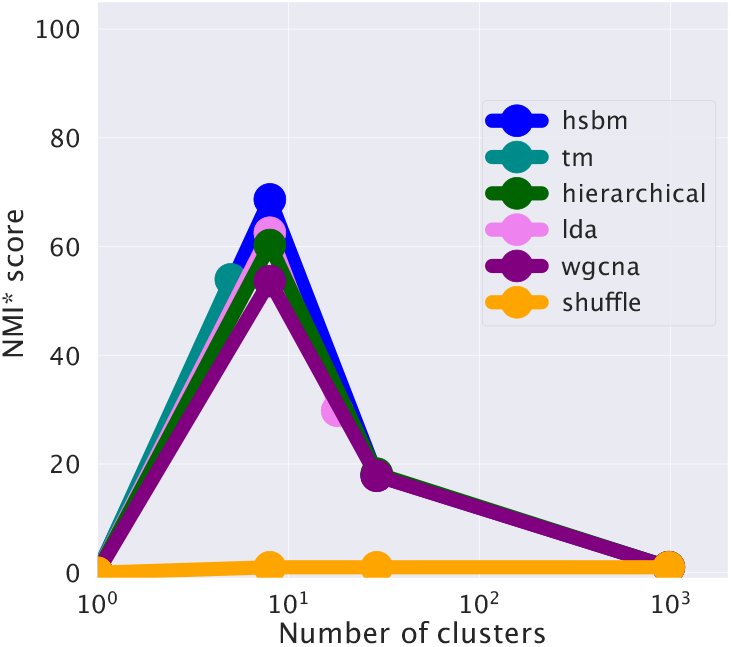
The *NMI* score of different algorithms. This score measures the agreement between the clusters obtained with the different algorithms and the ground truth sample partitioning defined by the tissues. All the algorithms reach a maximum score of around 10 clusters, which coincides with the number of tissues that we selected for the test. Topic modeling methods typically have the highest score.

Fig. 2 shows that at the first hierarchical level, corresponding to tissue partitioning, all the algorithms show rather good performance values, with a NMI* in the [50-70] range. hSBM and LDA perform slightly better than the other algorithms, but the performance gain is not significant on this simple clustering task. TM seems unable to recover the hierarchical organization of the data, probably due to its more rigid structure, and stops at the first partition layer. While at the level of sample clustering, all the algorithms behave in a quite similar way, their substantial differences manifest in the definition of topics, as the next section explains.

### 2.2 The effect of model assumptions and priors on the structure of the latent space

Topic modeling can also be interpreted as a projection of the data on a lower dimensional space in which the degrees of freedom are the topics. In this section, we investigate the structure of this latent low-dimensional space. In particular, we prove that different algorithms represent the data in latent spaces with very different properties, as a consequence of their different priors and inference procedures.

The structure of the topic space is defined by the topic distribution associated to samples. Specifically, the distribution *P* (topic|sample) for a sample can be considered as its coordinates in the low-dimensional topic space. We can then average over the samples belonging to a tissue to obtain *P* (topic|tissue) = Σ _sample∈tissue_ *P* (topic|sample), i.e., the projection of a whole tissue in the topic space. Since a topic is nothing but an ordered list of genes, these probabilities define which genes play a relevant role in identifying the tissues and defining the structure of the topic space. Building a dataset composed by samples from different tissues, thus with a well known structure, should help the biological interpretation of topics and of the genes involved. Identifying samples from different tissues should not be challenging, given their distinct expression patterns. In fact, most algorithms perform comparably well on this task (Figure 2). Consequently, one might expect that the latent topic structure inferred by various methods would be roughly similar.

Figure 3 reports *P* (topic|tissue) across tissues for two different algorithms. Figure 3a indicates that hSBM tends to find a common distribution for all tissues. Samples from all tissues are described by a mixture of a few important topics (i.e., with high *P* (topic|sample)), and several more specific but less present ones (with low *P* (topic|sample)). The way the topics integrate on a global scale differentiates the tissues

**Figure 3:**
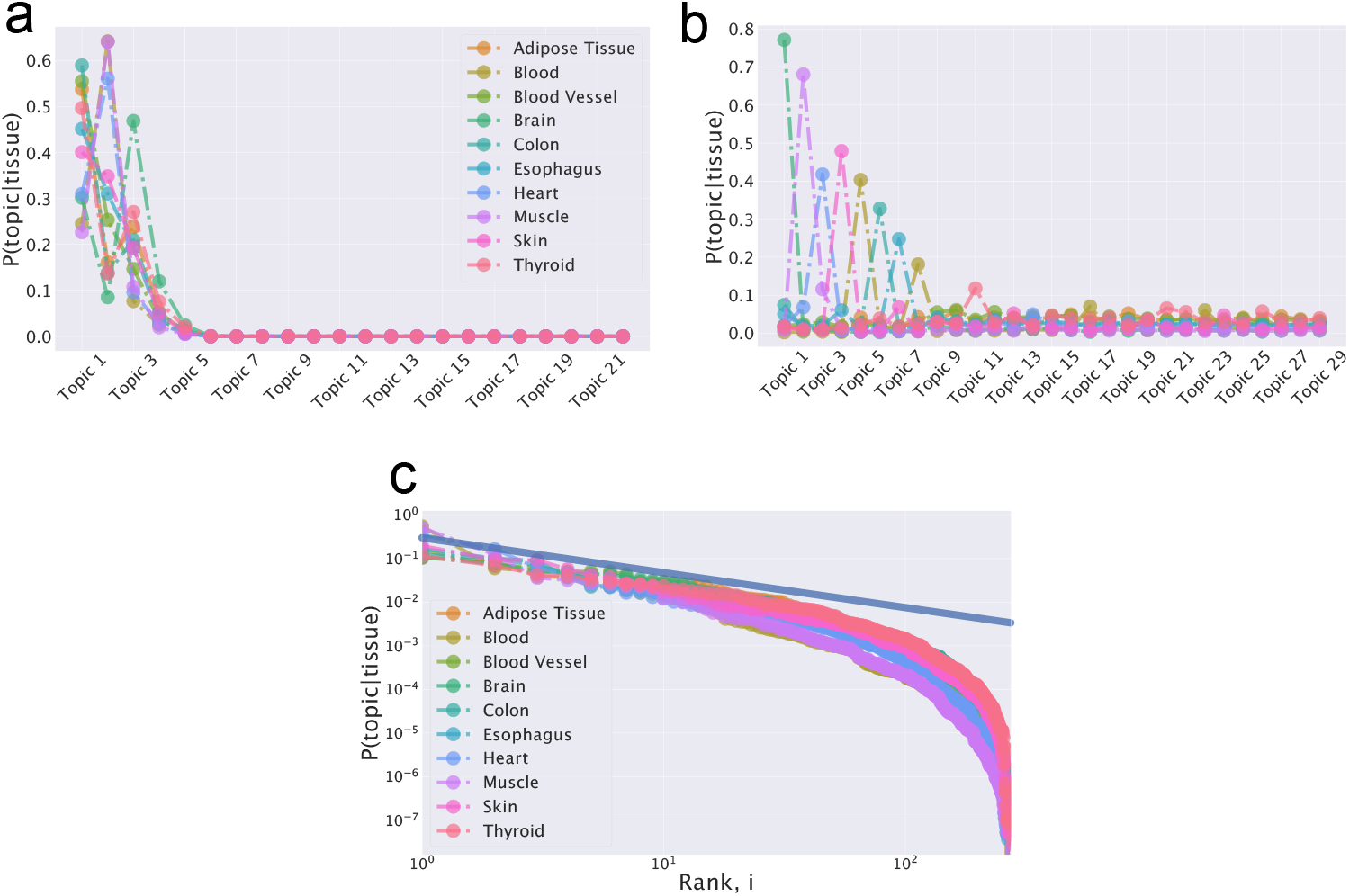
*P* (topic tissue) for different tissues. The topic importance in samples of different tissues is reported in **(a)** for the hierarchical Stochastic Block Model and in **(b)** for LDA using a dataset composed of samples from 10 tissues of GTEx and selecting the highly variable genes. The trend of the reconstructed topic space is affected by the algorithm priors. In **(c)**, topics are ranked in every tissue so that the first point corresponds to the most expressed topic in that tissue. For convenience, the plot is reported on a log-log scale. The power-law structure, at least for low ranks, of the original data space is preserved in the hSBM topic space. The analogous plot for LDA would be trivially peaked due to the probability distribution induced by the Dirichlet prior.

On the contrary, LDA leads to a radically different result due to its specific priors. The Dirichlet prior on which LDA is based forces the topic distribution towards more peaked solutions [5], as Figure 3b clearly shows in our case. As a consequence, the identified topic are much more tissue specific, with no common highly represented topics. This result is particularly appealing in the biological context because it potentially offer an easy interpretation. A sample is often characterized by just one or a few topics, and thus by a relatively small set of associated genes Nevertheless, this behavior is likely the result of the Dirichlet prior and, thus, the one-to-one association between topic and structure could not be actually present in the data. Similarly to LDA, Topic Mapping and WGCNA output a single layer of topics, with a structure of the topic space composed by a set of peaked distributions (see Supplementary Figure S1).

Gene expression levels are typically power-law distributed in most datasets [11, 25], and indeed a power-law distribution also well describes the expression levels in all the tissues (Figure S2 in the Supplementary Material). hSBM reflects this statistical property by reproducing an analogous distribution for the inferred topic structure (Figure 3c). This model was indeed originally proposed to abandon the specific LDA priors in the linguistic context, where power-law word abundance distributions are also ubiquitous, and was then applied to RNA-sequencing data with similar motivations [55, 56]. Here, we directly show the effect of this change of prior on the inferred topic structure.

The main difference in the structure of the inferred latent spaces can be observed using correlation analysis. We can measure the correlation between the typical expression of a gene in a topic ⟨*f*_*i*_⟩_*t*_ and the topic frequencies *P* (topic) in the samples. ⟨*f*_*i*_⟩_*t*_ is estimated by averaging the normalized expression of genes in each given topic, while *P* (topic) is the normalized abundance of topics in all samples. These two quantities are clearly correlated using hSBM, while this is not the case for LDA or other methods (Figure 4). In other words, hSBM finds the tissue structure in the data by defining topics that typically contain genes with similar expression levels. This is not true for other methods. Again, this is a consequence of the different model priors, and in particular of the uniform prior implicitly present in hSBM. Which topic organization is more desirable can be debated, and it is probably context-dependent. However, it is important to keep in mind that the specific gene grouping is strongly influenced by the selected method, especially when attributing a biological interpretation to the topics. In general, hSBM does no push towards few cluster-specific gene “markers”, but it rather organizes genes with similar expression values into topics and then uses the specific global combination of topics to cluster the samples. On the other hand, methods such as LDA tend to find topics composed by few genes with variable expression levels, and cluster of samples strongly associated to just one or few of these topics. While this is particularly appealing in terms of biological interpretation and subsequent experimental testing, it can be an artifact due to the model prior.

**Figure 4:**
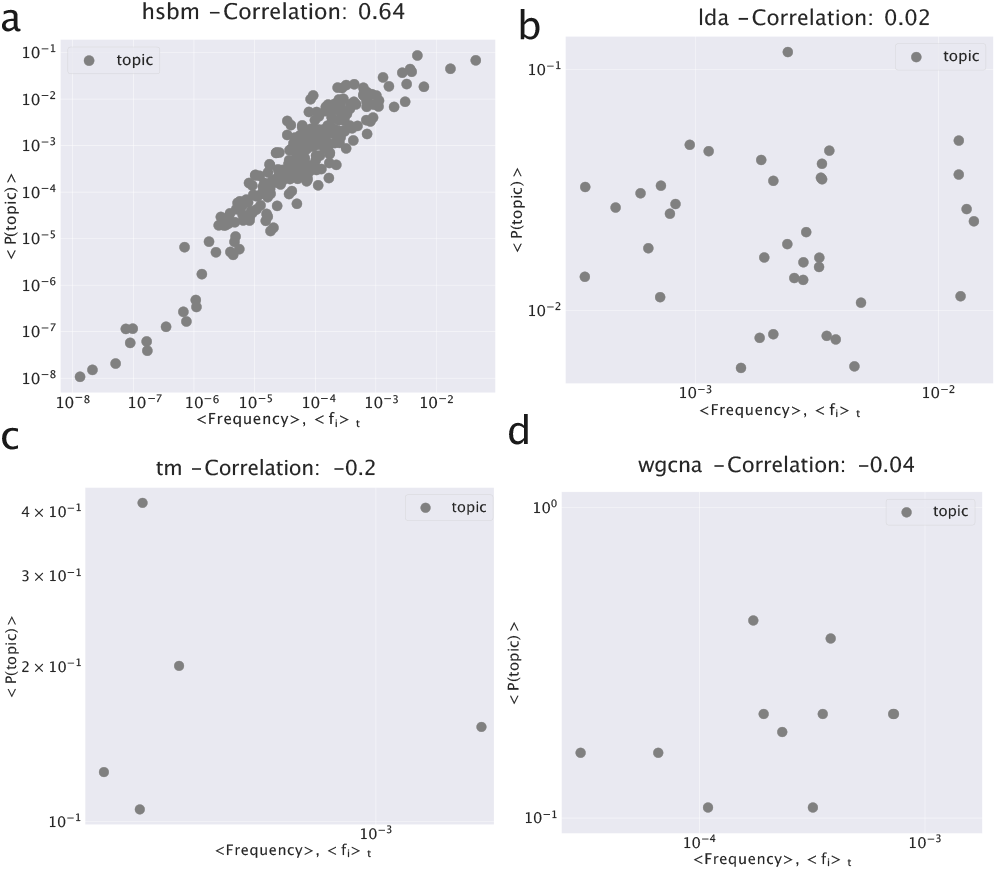
The average frequency of genes in each topic. We report the average normalized expression of genes in a topic: 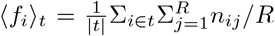, where *R* is the number of samples and *t* is a topic with |*t*| genes. We evaluate the correlation of this quantity with the frequency of topics in samples 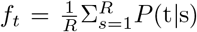. The different algorithms **(a)** hSBM, **(b)** LDA, **(c)** Topic Mapping and **(d)** WGCNA lead to very different correlation values. The topic distribution of hSBM is strongly correlated to the word distribution in the original data. This result resonates with a previous observation [12] about the widely different “disseminations” of topics when using hSBM on texts. This result can be reproduced at any level of the output hierarchy. We report here the hierarchical level of each algorithm with a number of clusters close to the number of tissues in the dataset (10), and a sufficient number of points to estimate the correlations: hSBM has 273 topics and 8 clusters, LDA 38 topics and 39 clusters. TM and WGCNA are not hierarchical models on the topic side, so we cannot select the number of topics.

The impact of the different priors becomes even more evident when the analysis is restricted to housekeeping genes. These genes are, by definition, consistently expressed in every sample. Their global expression pattern can still have enough information to separate different tissues, but we do not expect to find housekeeping “markers” of a tissue, i.e., housekeeping genes that are clearly differentially expressed in a specific tissue. Indeed, as expected, the distribution of hSBM topics is again power-law-like, and the obtained tissue separation is still quite accurate^1^, but based on global differences in topic composition (Supplementary Figure S3). On the other hand, LDA identifies single over-represented topics in different tissues in this case as well, likely due to the influence of the Dirichlet prior.(Supplementary Figure S3).

Biological processes result from complex interactions among potentially hundreds or thousands of genes. While reducing this complexity to a short list of “markers” may be appealing, it can be overly simplistic and may overlook genes that, acting in synergy with those markers, play a causal role in the process. Such oversight can hinder the development of effective therapies by missing potential targets. [51].

### 2.3 The relations between different tissues in the latent space

Since topic modeling can be viewed as a form of dimensionality reduction, a relevant question is how much of the structure from the original space is preserved in the low dimensional topic space. Our benchmark dataset is well suited to investigate this question. In fact, as reported originally by [30], there is a natural hierarchy of distances between tissues that can be observed at the level of gene expression patterns. We are interested in testing how much this hierarchy is preserved in the topic space created by different topic modeling techniques, in particular LDA and hSBM. Firstly, we define an “archetype” per tissue by averaging the *P* (sample|topic) over all its available samples. We use these prototypical points to measure the distances between tissues in the topic space. Analogously, the same calculation can be performed in the original expression space. We can thus associate to each tissue two alternative vectors of distances from all the other tissues, one evaluated in the original data space, and one in the topic space. At this point, the Spearman correlation between these two vectors provides information about the conservation between the tissue relations. The correlation is 1 if the ranking of the distances is perfectly consistent after the projection in the topic space. As shown in Figure 5a, the low dimensional projection of hSBM well conserves the tissue hierarchy in the expression space. Interestingly, this is only partially true for LDA (Figure 5b), while with Topic Mapping (Figure 5c)) and WGCNA (Figure 5d) the tissue structure appears to be lost.^2^

**Figure 5:**
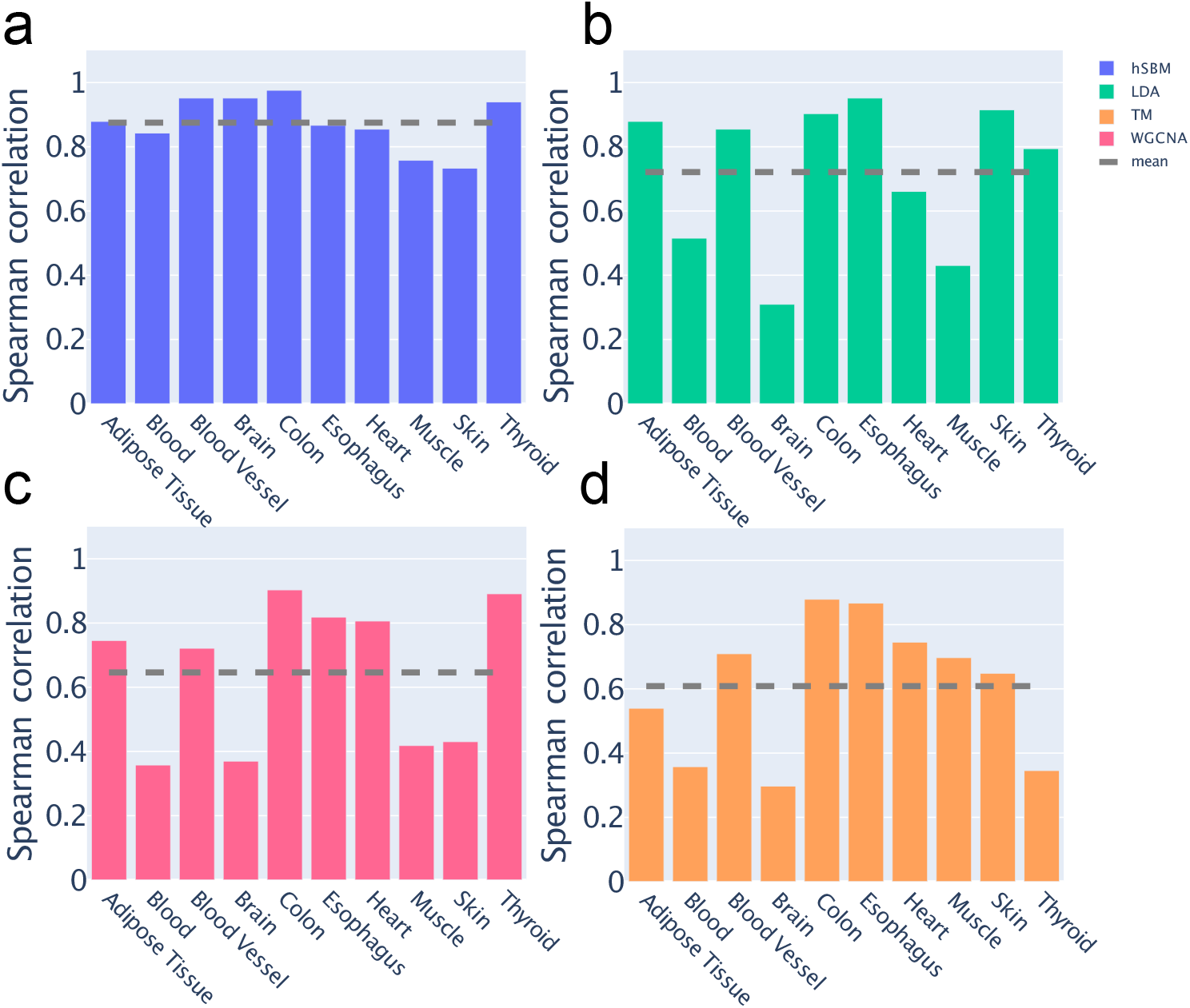
Spearman correlation between vectors of tissue distances calculated in the expression space or in the topic space. A comparison between different algorithms (**a**) hSBM, (**b**) LDA, (**c**) Topic Mapping, (**d**) WGCNA is reported. The dashed lines correspond to the mean value across tissues. High correlation values indicate a conservation of the tissue relations in the topic space.

### 2.4 Different methods recover similar clusters but different topics

The phenotypic differences observed across tissues are strongly reflected in their gene expression profiles. In fact, nearly every algorithm tested was able to effectively distinguish between samples from different tissues. This is confirmed by the high Normalised Mutual Information (NMI) [44] (see Methods Section for a detailed discussion of the model evaluation measures) values obtained, using the known tissue structure as ground truth (Figure 2). Indeed, different algorithms converge to very similar partitions at the level of samples, at least at the coarse resolution of tissues (Figure 6a). However, as discussed in the previous sections, the topic spaces generated by alternative models exhibit structural differences. We now directly assess the overlap in the genes that the various models prioritize to achieve tissue separation.

**Figure 6:**
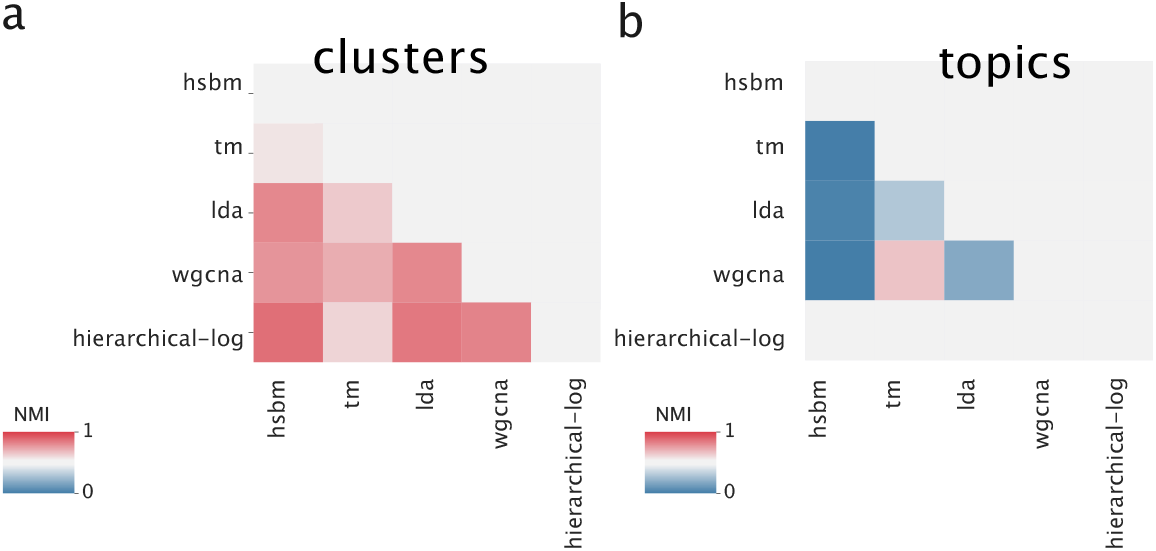
Similarity between the partitions obtained with different algorithms on the sample side (a) and on the gene side (b). The similarity is evaluated using the NMI* score, as defined in the Methods section. Different algorithms build very similar sample partitions but tend to build non-overlapping topics. Hierarchical clustering does not provide a topic structure, therefore it cannot be compared with other methods at the gene level. Most of the algorithms present a hierarchical structure, in this case, we compared them at a single point of resolution. We choose a resolution in which all algorithms present a compatible number of topics (hSBM 22, LDA 8, TM 5, WGCNA 21) and clusters (hSBM 29, LDA 28, TM 5, WGCNA 30, hierarchical 29).

Figure 6b reports the agreement between the partitions of genes obtained with different models. The low NMI scores show that the overlap between the gene clusters is quite low. Therefore, the different model priors not only lead to distinct structures of the topic space but also produce varying associations between genes and topics. This implies that the gene sets linked to specific tissues are highly model-dependent.

In general, machine learning methods, such as topic modeling, leverage statistical associations to perform a task without any domain knowledge. These statistical associations obviously do not necessarily correspond to causal relations [13, 42, 59]. Given that the number of genes often exceeds the number of samples and that gene expression levels are frequently correlated (e.g., genes regulated by the same transcription factor), multiple combinations of genes could effectively correlate with the tissue structure. Consequently, it is not surprising that different models, each with distinct priors, may focus on different gene sets for clustering the samples. However, the substantial differences in the gene sets associated with the samples serve as a caution to users: these associations should not be overinterpreted, and the biases inherent in each model should be carefully evaluated.

### 2.5 Gene Sets Enrichment Analysis confirms the biological relevance of the topics

Topics are essentially lists of genes and thus gene set enrichment analyses can be used to identify the biological features associated to topics. We start this analysis from the tissue structure identified by hSBM. We first build a centered version 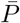 (topic|tissue) of the probability distribution *P* (topic|tissue) by subtracting the mean probability value. This centered version highlights the topic that are over-represented in a given tissue as displayed in Figure 7.

**Figure 7:**
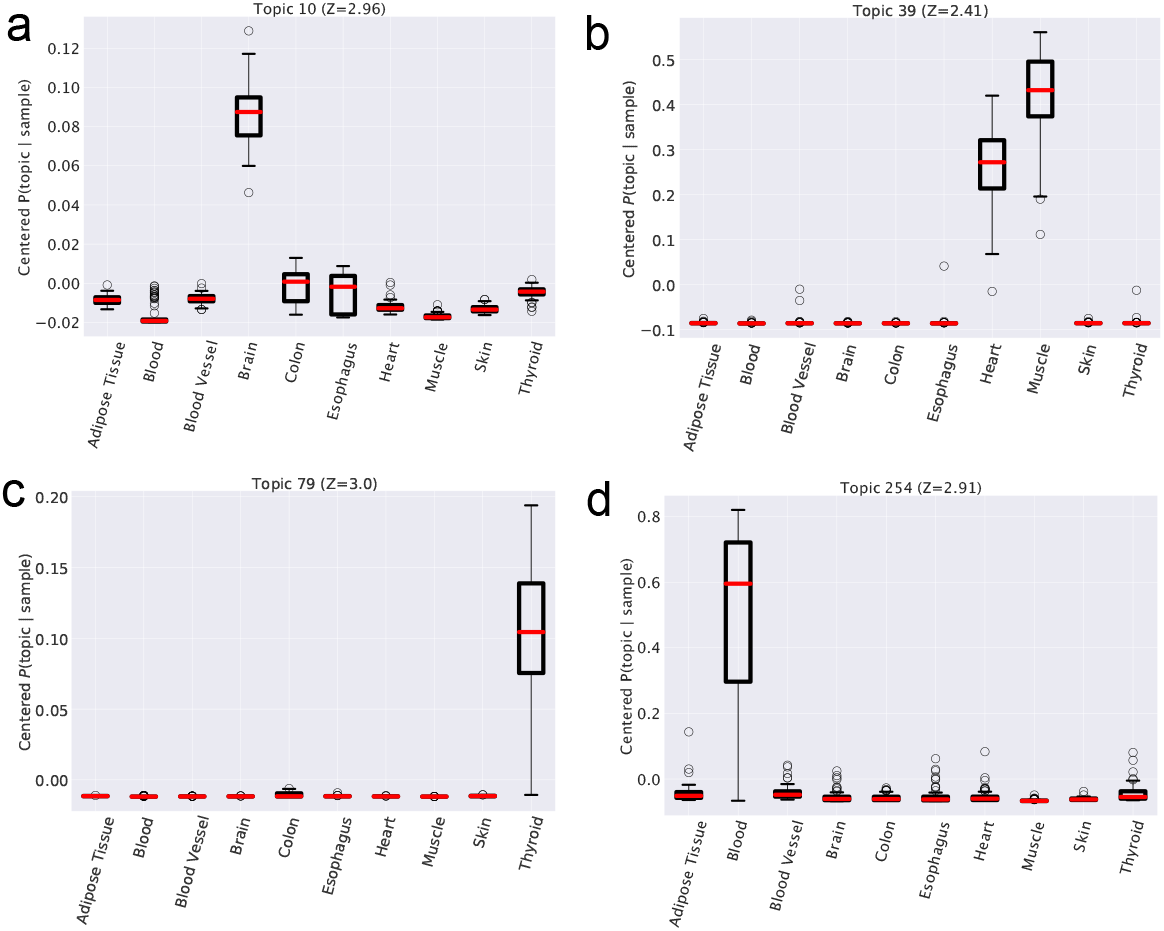
Distribution of centered 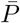 (topic|sample). We grouped 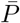 (topic|sample) for all samples belonging to the same tissue in each box. Each subpanel refers to a specific topic. This value reveals, in the probability distributions, trend correlated with the tissues. We reported in this panel some of the more informative topics. In Table 1 we reported gene ontologies of genes in these topics.

We can now isolate the topics whose centered probability is higher in one or more tissues using a z-score to evaluate the significance of these deviations. Finally, we can apply a Gene Set Enrichment Analysis [53] to the gene lists associated the significant topics. An analogous procedure was previously used to interpret the biological meaning of topics in the context of cancer transcriptomics [55]. Illustrative examples of this analysis are reported in Figure 7 and in Table 1. In many cases, a single topic is over-represented in a tissue and thus should be representative of the tissue biological properties. This is the case of topic 10 in Fig. 7a, which is strongly associated to brain tissues. The genes in the topic are indeed enriched in neuronal GO categories (see the first entry of tab.1). In some other cases, a single topic is enriched in more than one tissue, probably due to common biological pathways or functions in the two tissues. This is the case of topic 39, which is associated both to Muscle and Hearth (Fig. 7b), and accordingly to the keywords “myosine”, “myogenesis” and “muscle contraction”. Other examples are reported in Figure 7 with the corresponding enrichment results in Table 1.

**Table 1:**
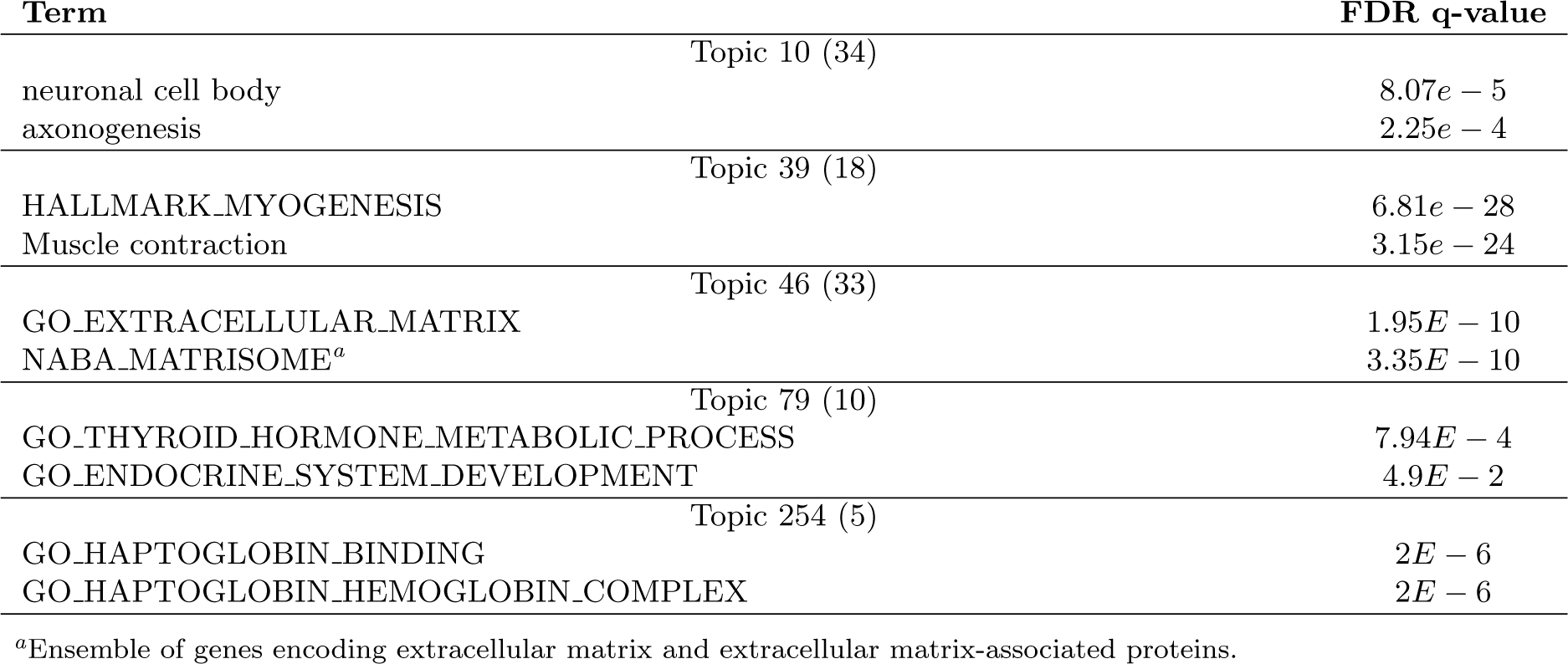
Gene ontology enrichment tests performed on the gene lists (number of genes reported in brackets) associated to different topics using GSEA [53].

| Term | FDR q-value |
| --- | --- |
| Topic 10 (34) |  |
| neuronal cell body | $8.07e - 5$ |
| axonogenesis | $2.25e - 4$ |
| Topic 39 (18) |  |
| HALLMARK_MYOGENESIS | $6.81e - 28$ |
| Muscle contraction | $3.15e - 24$ |
| Topic 46 (33) |  |
| GO_EXTRACELLULAR_MATRIX | $1.95E - 10$ |
| NABA_MATRISOME<sup>a</sup> | $3.35E - 10$ |
| Topic 79 (10) |  |
| GO_THYROID_HORMONE_METABOLIC_PROCESS | $7.94E - 4$ |
| GO_ENDOCRINE_SYSTEM_DEVELOPMENT | $4.9E - 2$ |
| Topic 254 (5) |  |
| GO_HAPTOGLOBIN_BINDING | $2E - 6$ |
| GO_HAPTOGLOBIN_HEMOGLOBIN_COMPLEX | $2E - 6$ |

<sup>a</sup>Ensemble of genes encoding extracellular matrix and extracellular matrix-associated proteins.

Interestingly, some topics are common to several (or all) tissues. In the context of natural language processing, these topics have been characterized as “common topics” containing less informative words, such as articles [12], which are present in all texts. Analogously, in our context, these common topics are enriched in general GO terms, which are shared by all the tissues. This is the case, for instance, of topic 46 (see the third line of Table 1 which is enriched in keywords like “extracellular matrix” which can be associated to different tissues.

We performed the same functional analysis for the topics identified by the other three algorithms. All the results are presented in Figures S5 (LDA), S6 (TM), and S7 (WGCNA) of the Supplementary material. The GO enrichment is relatively strong for WCGNA and TM topics, while slightly weaker for LDA ones. LDA does not naturally define the topics as gene lists, instead its output is a probability distribution whose support is the whole gene repertoire. As discussed in the Methods section, we select the 20 genes with the higher associated probability given a topic (20 the typical number of genes per topic found by hSBM).

An issue related to the WCGNA results is that the gene list associated to a topic is in general very large (one order of magnitude larger than the avrage hSBM list) making the topics much less specific. On the contrary, TM outputs on this dataset a small (i.e., 5) number of topics to identify a one-to-one relation between topics and tissues. Possibly, these problems could be made less severe with a suitable choice of hyperparameters. However, as we mentioned in the introduction, in order to make a fair comparison with paramter-free methods such as hsbm and to avoid overfitting we did not perform model-specific explorations of hyperparameter values, and adopted the standard settings.

Interestingly, even if the topics found by different methods are very different, as shown in the previous sections, the associated gene lists often shows significant enrichment for gene ontologies that are compatible with the tissue structure (see Supplementary Tables S5, S6 and S7).

### 2.6 The use of topic modeling to distinguish healthy from cancerous tissues

As an additional relevant benchmark dataset, we combined samples from healthy tissues from GTEx [30] and cancerous tissues from TCGA [57]. As a consequence, two main strucures are present in this dataset: the tissue structure and the distinction between healthy and cancer tissues.

The results of applying hSBM and LDA to this dataset are reported in Fig 8a and Fig. 8b, respectively. We display the NMI* levels evaluated with respect to the tissue structure only (blue lines), or to the tissue structure with the additional partition due to status, i.e., healthy vs cancerous (red lines). The task of retrieving even just the tissue structure is now more challenging due to the additional variablity introduced by the presence of cancerous samples. Indeed, the NMI* values are lower than those reported in Figure 2. In this more complex setting, hSBM outperforms LDA in discovering both structures, further confirming its potential in the context of cancer transcriptomics [55].

**Figure 8:**
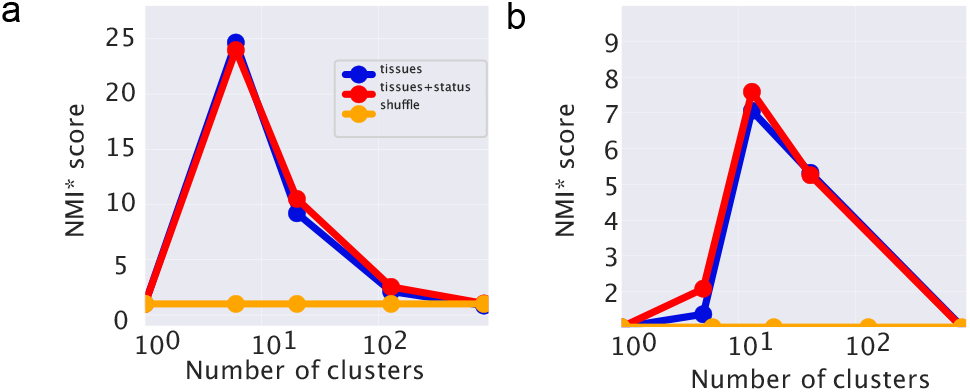
Identifying healthy and cancer tissues. We report the NMI* scores (see the Methods section) for hSBM **(a)** and for LDA **(b)** when applied to a dataset combining healthy and cancerous samples from 20 different tissues. The score is evaluated with respect to the tissue structure (blue lines) or concerning the tissue structure with the additional status annotation (red lines).

The lower resolution level is dominated by the tissue organization. In fact, hSBM typically finds a number of clusters close to the number of tissues (10) and the NMI is higher if evaluated with respect to the tissue structure. At the subsequent layer, the algorithm further separates reasonably well the healthy tissues from the cancerous ones. Indeed, the red line (referring to the partitioning with health status) becomes higher than the blue line, where the number of clusters is generally close to 20 as expected (Fig 8a). This effect is less evident using LDA (Fig 8b).

### 2.7 Robustness to data pre-processing and gene selection

Data pre-processing and gene selection procedures to reduce the number of features are standard practices in the analysis of RNA-sequencing data [52]. However, most protocols do not have a clear theoretical motivation and can potentially affect downstream analysis and conclusions. In this section, we focus on the often-used data logarithmic transformation and on gene selection procedures to analyze their effect on the results of topic modeling algorithms.

#### 2.7.1 Topic modeling algorithms do not need data log-transformation

The mRNA counts from RNA-sequencing experiments are the result of a sampling process, thus their variability is due to a combination of biological and technical sources [45, 25]. For this reason, the data are typically first normalized to reduce the sampling effect. Moreover, the data are often log-transformed [6] with the underlying hypothesis that fold-change variations are more relevant than absolute changes. In fact, the euclidean distance of sample *j* from sample *j*′ is defined as 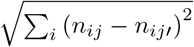, but it becomes 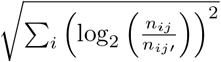 with the log-transformation, making the relative fold-change 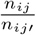, the only relevant quantity. By affecting how the distances are evaluated, this pre-processing procedure can affect the results of clustering methods. Therefore, we want to compare the performances of different algorithms using samples in the original data space (using the standard Transcripts Per Million or TPM values) or samples in the log-transformed space. Figure 9 summarizes this comparison. It is clear that topic modeling methods, both hSBM and LDA, do not need this pre-processing step to obtain significant scores in the task of finding the tissue structure in the dataset. On the other hand, there is a relevant drop in performance for both WCGNA and hierarchical clustering if the log-transformation is not applied. Topic Mapping was not included in this analysis because it considers a network of genes based on co-occurences that are unaffected by a log-transformation. A gene is present in a sample if TPM *>* 0 and this is equal to log_2_(TPM + 1) *>* 0.

**Figure 9:**
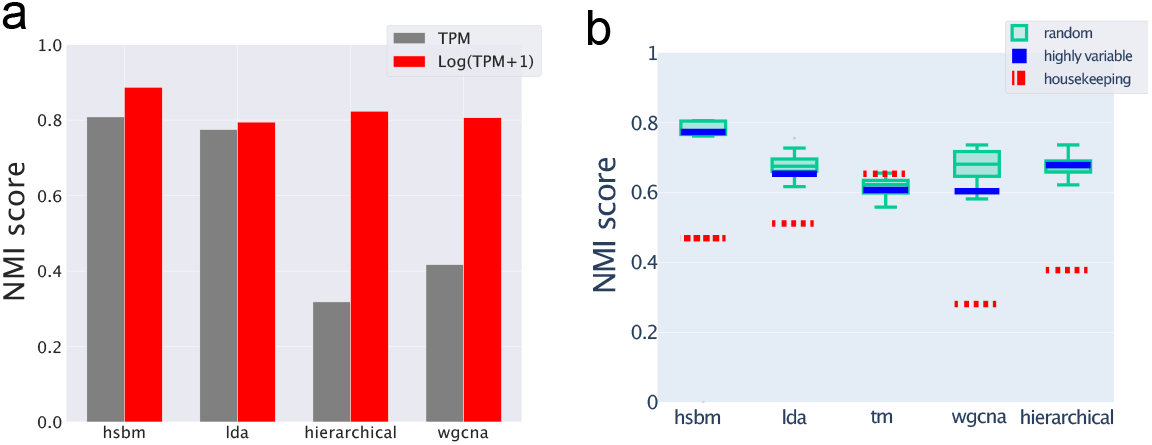
Robustness to data pre-processing. **(a)** The performances in terms of NMI of different algorithms in retrieving the tissue structure when operating on *TPM* expression values (grey) or on log-transformed data using log(*TPM* +1) (red). **(b)** Comparing different gene selections. We reported the NMI score of the hierarchy level with the number of clusters more similar to 10 (the real and expected one) (for some algorithms, e.g. LDA, it is not possible to force the number of clusters, that’s why it may vary. See Method sections for details). The red dashed lines represent the particular choice of 3 000 housekeeping genes from ref. [9]. The scores obtained using the standard selection of highly variable genes are reported as blue lines. The green boxes report the scores for different algorithms when choosing a random subset of genes as features (for each selection and each algorithm we chose the score where the algorithm found a number of clusters more similar to 10). We use directly the NMI, without the normalization with respect to a random partition, since the number of clusters is quite constant in this analysis.

The log-transformation has several potential pitfalls. For example, the zero count values have to be replaced by a small constant before the transformation, and this can introduce artifacts in sparse datasets. Moreover, the log-transformation compresses differences at higher expression levels that could be biologically relevant. Therefore, it is important to know which algorithms can actually operate reliably in both expression spaces and our analysis shows that this is the case for topic modeling methods.

#### 2.7.2 Robustness to gene selection

Another standard procedure in the analysis of RNA-sequencing data is gene selction [46]. Reducing the number of features is often necessary to speed up downstream analysis, and can also potentially eliminate less informative features that can act as confounding factors. This section discusses the robustness of different algorithms with respect to feature selection by considering the two extreme cases of random gene selection and of selection of housekeeping genes. While typically an arbitrary number of highly variable genes is used (and this is the selection criterium we adopted so far), these more radical criteria can better highlight the algorithms’ dependences on gene selection.

Actually, the ability of different algorithms in retrieving the tissue structure when using highly-variable genes or a random selection of genes is not significantly different (Figure 9b). This result suggests that the information about the different tissue phenotype is redundantly present in the expression profiles. Therefore, many different subsets of features can be used without significant differences.

Housekeeping genes are instead expressed in nearly every sample and typically exhibit high expression levels and low variability. Therefore, by selecting only housekeeping genes we are defining the more difficult task of retrieving the tissue structure using small and collective differences in otherwise quite conserved features. Figure 9b shows that topic modeling algorithms, can still reasonably well reconstruct the tissue structure in this setting (NMI scores as red dashed lines), while standard clustering algorithms, such as WGCNA or hierarchical clustering, have a more evident performance drop.

### 2.8 Sample classification in the latent space

Topic models can also be interpreted as dimensionality reduction techniques. In fact, they define an embedding space (i.e., the topics) in which the data can be projected. A simple Neural Network (NN) can be trained in a supervised manner to perform, for example, healthy vs tumor classification of samples represented in this embedding space. The idea is to use the unsupervised learning of a topic model to learn suitable sample representations, and then build a classifier in this representation space to annotate new samples. This procedure is analogous to the unsupervised pre-training that has been adopted in several deep learning projects [3].

In practice, each sample is only defined by a vector of its topics, reducing the ≃ 20 000 gene features to ≃ 1800 topics. The number of topics depends on the selected hierarchical level of hSBM and sets the dimension of the input space for the NN. We can thus use a simpler and faster-to-train model in the topic space without losing explainability since, as shown in the previous sections, topics can be associatedto biological functions with gene set enrichment analysis.

Using the topic space defined by hSBM, we trained a NN for two classification tasks: tissue separation and healthy vs cancerous sample. A standard cross-validation setup has been adopted, as presented in the cartoon of Figure 10a. We also compared our results with a simpler K-Nearest Neighbour (K-NN) classifier that can assign the labels using the sample position in the original expression space.

**Figure 10:**
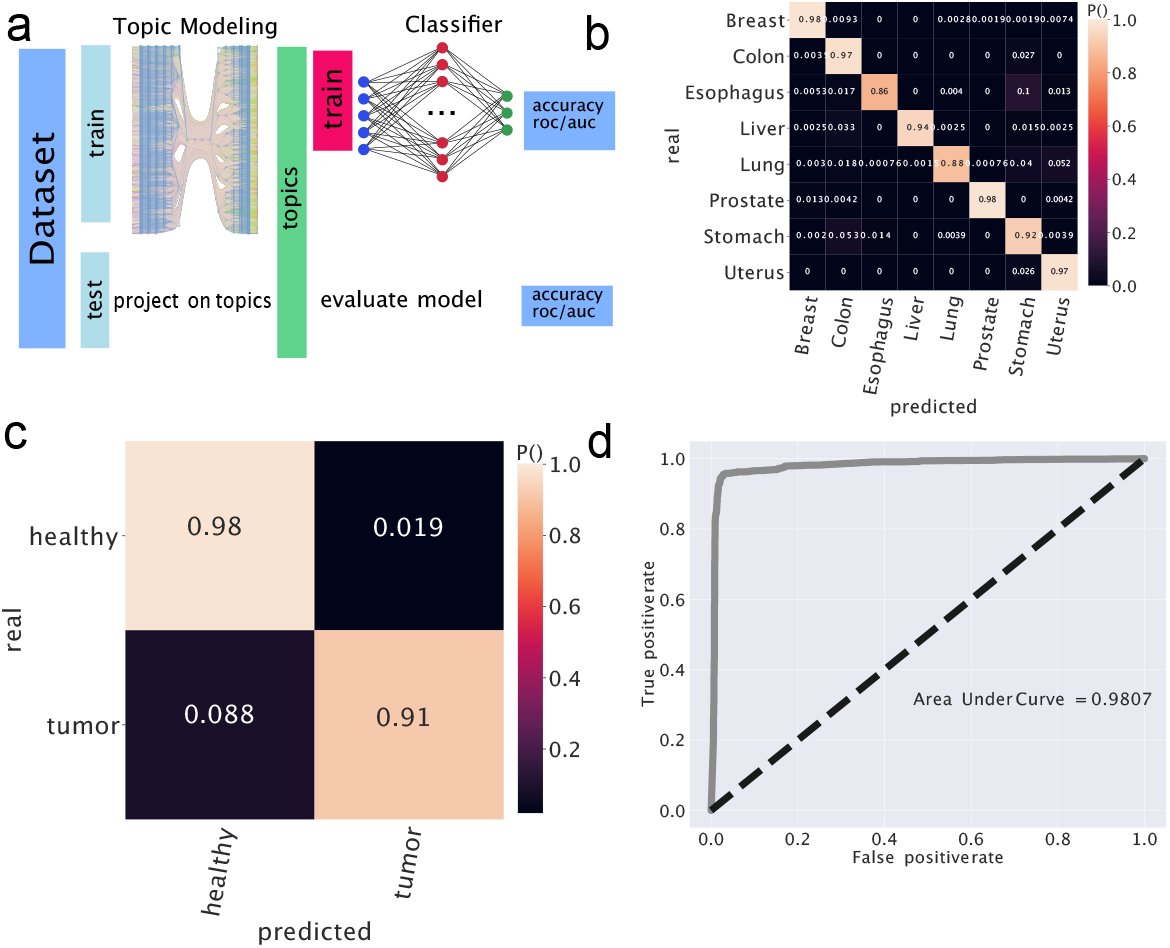
Prediction on the topic space. **(a)** Topology of the experiment. **(b)** The confusion matrix of the experiment involving tissues. **(c)** We run a similar experiment but try to predict if a tissue is healthy or diseased. The setting was the same as the tissue except for the tuning of some hyper-parameters and the fact that the last layer uses sigmoid as an activation function. In the figure the confusion matrix. In **(d)** the ROC curve with the thresholds. We considered *positive* healthy samples.

The first task is to classify the tissue origin of new samples (not previously used to define the topic space) by first projecting them in the topic embedding space. A small NN trained in this space obtains an accuracy of 0.93 on this task. Figure 10b reports the confusion matrix for this setting. Since the task is not particularly complex, a K-NN can be used to assign tissue labels with an accuracy around 0.8 in the original data space, and around 0.9 in the log-transformed expression space. Similar performance values can be obtained by training the NN in the original expression space (the full performance metrics are reported in Tables S3 and S4). However, the advantage of the projection in the low-dimensional topic space is that we can train much smaller models with similar performance values and interpretable results. Moreover, as the size of expression datasets increases, the information extracted by the unsupervised pre-training with topic modeling is expected to continuously increase the performance on complex downstream tasks, even if defined by very small annotated datasets for supervised training.

The second task we tested was the ability to classify healthy vs cancerous samples. A NN operating in the topic embedding space obtains an accuracy of 0.95 and an Area Under Curve score of 0.98. The confusion matrix and the ROC associated to this binary classification problem are reported in Figure 10c,d.

This analysis is a proof of concept that a simple neural network can be used successfully in the structurally rich but low dimensional topic space for different downstream classification tasks.

### 2.9 Conclusions

Topic modeling is a simple yet powerful unsupervised learning technique that can be naturally applied to transcriptomic data, leveraging on the strong statistical analogies with datasets of natural language [55, 25]. While few applications in this context have been explored [8, 55, 56, 61], we are still at the early stages of understanding its full capabilities a possible pitfalls. By comparing different topic models and standard clustering algorithms on a benchmark dataset with a partially known internal structure, we demonstrated that topic modeling can cluster expression samples as effectively as traditional methods. However, topic models offer a richer probabilistic framework, allowing for the extraction of latent variables and their associations with genes in a probabilistic manner. This framework also provides a graded sample membership rather than a mere grouping. Moreover, topic models appear to be generally robust with respect to data preprocessing and feature selection and can be used as a dimensionality reduction method.

Each topic model employs different priors. We have shown how these priors strongly influence the topics identified by the methods. This is especially relevant for transcriptomic data, in which relatively few samples are embedded in a very high-dimensional expression spaces. As a consequence, there are potentially many unconstrained degrees of freedom, making the choice of statistical priors particularly important. The presented results underscore the need to consider these priors carefully when selecting a topic modeling approach, as they shape the model focus and its ability to uncover biologically relevant structures.

Among the tested topic models, the hierarchical Stochastic Block Model (hSBM) offers several distinct advantages. It does not require adjustable parameters by automatically detecting the number of topics, and naturally provides a hierarchical structure of the output. Moreover, our results demonstrate that hSBM captures the ubiquitous power-law distribution of gene expression values, projecting it onto an analogous topic distribution. Additionally, hSBM maintains the hierarchical relationships between hidden structures (the tissue structure in our case) in the embedding space. These features make it particularly well-suited for capturing potentially complex structural relations in transcriptomic data.

Despite its advantages, hSBM has some limitations. In particular, it relies on time-consuming Monte Carlo simulations, making it slower compared to models like Latent Dirichlet Allocation (LDA). LDA, with its Dirichlet prior, may be particularly suitable for identifying short lists of “marker” genes due to its tendency to identify single, over-represented topics within sample clusters. In contrast, hSBM naturally emphasizes global rearrangements of expression patterns, likely due to its “flat” prior. Therefore, using hSBM and LDA in tandem could offer complementary insights, with LDA supporting marker extraction and hSBM providing a broader view of the overall structure.

Given the field of genomics often focuses on a limited set of well-studied genes [51], it is important to include algorithms like hSBM in the analytical toolbox to capture global patterns and structural alterations that can distinguish between samples.

Topic modeling is still an active research area. In particular, there are recent developments of mathematical descriptions based on the stochastic block model that are still waiting to be adapted and tested for genomic data. For instance, merge-split Markov Chains [39] or Bayesian nonparametric formulations [62] could lead to improvements in the performance of hSBM-like algorithms. Moreover, in this work, we ran hSBM with multiple initializations and choose the one with the shortest description length. However, the recently proposed technique based on consensus [40] could be an interesting alternative to explore.

## 3 Materials and Methods

### 3.1 The Genotype Tissue Expression dataset

The Genotype-Tissue Expression (GTEx) Project was supported by the Common Fund of the Office of the Director of the National Institutes of Health, and by NCI, NHGRI, NHLBI, NIDA, NIMH, and NINDS. The data used for the analyses described in this manuscript were obtained from the GTEx Portal phs000424.v8.p2 [26]. GTEx data were downloaded in Transcript Per Million (TPM) format.

We also downloaded from the GTEx portal https://gtexportal.org the annotations of samples and in particular we focused on the tissue type (the area from which the tissue sample was taken) and its subtypes (SMTS and SMTSD labels).

We selected 1 000 samples from each one of the 10 most represented (in terms of number of samples per tissue) tissues: Adipose Tissue, Blood, Blood Vessel, Brain, Colon, Esophagus, Hearth, Muscle, Skin and Thyroid.

### 3.2 Algorithms

#### 3.2.1 hierarchical Stochastic Block Model

We used the stochastic block modeling topic modeling `sbmtm` class in the code we forked from [12] available at https://github.com/martingerlach/hSBM_Topicmodel/tree/develop. We passed the gene expression matrix as a weighted graph to the model.

The hierarchical Stochastic Block Model is a kind of generative model that tries to maximize the probability that the model describes the data

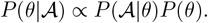

Its approach is completely non-parametric and aims to maximize the posterior probability. We used the `minimise_nested_blockmodel_dl` (the degree corrected version of the model was used) function from graph-tool [34] to minimize the description length Σ = − *lnP* (𝒜|*θ*) − *lnP* (*θ*) in a nested version of the model [35, 33, 36, 37, 38]. Minimising Σ leads to the maximization of the posterior probability. In our setting 𝒜 is nothing but the gene expression matrix, weighted by the number of *TPM* and *θ* refers to the model parameters. It is a two-dimensional matrix with entries 𝒜_*ij*_: 𝒜_*ij*_ is the expression value of gene *i* in sample *j*. In a setting where rows are words and columns are documents it is often referred to as “Bag of Words”. We performed model selection minimizing the description length Σ 10 times and choosing the model with the shortest description length.

#### 3.2.2 LDA

Specifically, we adopted the implementation provided by scikit-learn [32, 16]. We used the default setting for the parameters *α* and *β* (they were both set to 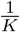, being *K* the number of topics). When managing LDA output, we selected the *argmax* of *P* (topic sample) to define clusters. Note that it is not guaranteed that every topic is maximized in at least one of the samples, so the number of clusters can be smaller than the number of topics. Using this procedure it is not possible to select a-priori the number of clusters. A similar procedure was used to identify clusters with Topic Mapping.

To get a list out of the LDA topic distribution we selected the 20 most distinctive genes of each topic. The distinctiveness was described and used in LDA analyses of GTEx by [8] and is the minimum Kullback-Leibler distance of *P* (topic|gene).

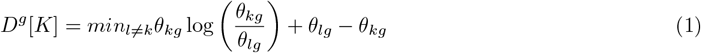

being *θ*_{*k,l*}*g*_ the *P* (gene_*g*_ |topic_{*k,l*}_).

#### 3.2.3 Topic Mapping

We ran Topic Mapping [23], with default parameters: the number of runs -r was set to 10 and the minimum topic size -t was set to 10 to avoid small useless clusters and force the algorithm to put at least 10 samples per cluster.

Topic Mapping requires as input a corpus of text and not a Bag of Words or a design matrix, this is why we built a corpus in which each text was constructed from the 1 000 most expressed genes in each sample. In other words, each sample was translated into a text composed of the 1 000 genes most represented in that sample.

Clusters are defined using the *argmax* of *P* (topic|sample).

Finally, we got the list of genes from the output of Topic Mapping setting a threshold on *P* (topic|gene) corresponding to the 99^*th*^ percentile and kept all the genes above that threshold.

#### 3.2.4 Weighted Gene Correlation Network Analysis

We considered the R implementation by https://cran.r-project.org/web/packages/WGCNA/. WGCNA automatically outputs sets of genes (the topics) and it automatically selects their number. We considered WGCNA [24, 63] modules as topics, in particular, we choose genes with a correlation higher than 0.5 to select the genes within the modules.

#### 3.2.5 Hierarchical clustering

We use the agglomerative clustering class implemented in *sklearn* and set to use *euclidean* affinity and *complete* `linkage` following [8]. We would like to recall that our purpose is to use as less a-priori knowledge as possible, that’s why, under reasonable assumptions, we choose standard settings without grid search hyper-parameters or applying other optimization methods. hSBM provides all the information we discussed without any a-priori assumption or parameter.

The input for the various algorithms was a matrix 𝒜 whose entries are the gene expressions. Data of this matrix are samples and features are genes. In literature, this is often called “Bag of Words”.

One of the advantages of hSBM is that it automatically outputs the number of clusters and the clusters themselves. hierarchical and WGCNA output a tree of samples; we need to cut it to obtain clusters. In LDA the number of topics is a parameter and in TM we set the minimal size of topics.

#### 3.2.6 Computational complexity

hSBM was run on a node of a cluster [1] with 48 cores and 700 GB of memory and it took several hours to be run, all the other algorithms take minutes on a laptop with 2 cores and 8 GB of memory. The complexity of hSBM is *O*(*V Ln*^2^*V*) only in the case of sparse networks when *E* ∼ *O*(*V*) [35], LDA has a complexity of *O*(*K* * *N* ^2^) where *N* is the number of genes, *V* the number of vertices (samples and genes), *E* the number of edges, and *K* the number of topics. In our setting we have *V* = 1000 + 3000 = 4 000 vertices, *E* ∼ 450 000 edges and tens of topics *K*. In this setting (i.e. dense network with 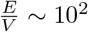) the complexity of hSBM is higher than the complexity of LDA.

### 3.3 Models evaluation

We evaluated the performance in clustering by looking at the partition of samples between clusters. We used the V-measure or Normalised Mutual Information *NMI* [44] to evaluate the performance of the model in the task of separating samples of different tissues. A similar approach involving *NMI* was proposed by [50] to evaluate topic modeling performance in reconstructing synthetic corpora. *NMI* is estimated as the harmonic average of homogeneity and completeness. Homogeneity is defined as 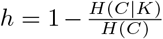 and completeness is 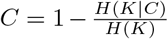, so 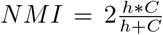. In our setting *C* is the ensemble of GTEx tissues and *K* is the partition in output. In, being the number particular 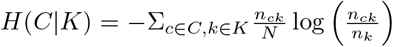 and 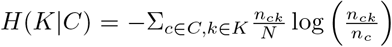 of samples of tissue *c, n*_*k*_ the number of samples in cluster *k* and *n*_*ck*_ the number of samples of tissue *c* in cluster *k*. If all the nodes in cluster *k* belong to tissue *c n*_*k*_ = *n*_*ck*_ and the homogeneity *h* = 1. Similarly, the completeness *C* equals 1 if all samples of tissue *c* are in the same cluster *k*. See Table S5 in the Supplementary for a brief illustration of these metrics.

We estimated the *NMI* score for all algorithms varying the resolution (layer).

The *NMI* is not zero in the case of random sampling: when the number of clusters is too high the result is almost random for every algorithm. That’s why we chose to divide the score by the one obtained with a random partition (we shuffled the data maintaining the number of clusters and the clusters sizes). The *NMI* for the random model was estimated for the number of clusters in output by hSBM, other points are interpolated from this. We referred to the *NMI* scaled to random as *NMI*^*^.

We applied this approach to TPM data and on transformations of them. In particular, we applied two different log-transformations. We estimated log_*b*_(*TPM*_*ij*_ + 1) being *TPM*_*ij*_ the number of Transcript Per Million of gene *i* in sample *j*. We tried with base *b* = 2 and *b* = 10 obtaining the same results.

### 3.4 Hyper-geometric test and model comparison

Results of different algorithms were compared using two different approaches.

First, given the list of genes in two topics of different algorithms, we performed an hyper-geometric test. The test consists in evaluating the probability that the overlap between the two sets of happens by chance

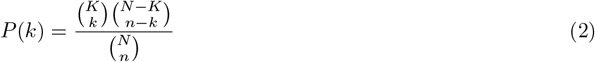

where *N*, the population size, is the number of genes processed by both algorithms. *K*, the number of successes in the population, is the number of genes in the topic of the first algorithm. *n*, the sample size, is the number of genes in the topic of the second algorithm. The number of successes *k* is the intersection between the two sets i.e. the number of genes in common between the topics obtained by the two algorithms.

Note that some algorithms drop genes during the analyses. WGCNA has the *goodSamplesGenes()* function that returns a list of samples and genes that pass some criteria: missing entries, entries with weights below a threshold, and zero-variance genes. When we built corpora for TM it was not guaranteed that every gene was picked up at least once, and the same for the LDA topic-construction procedure, some genes can be left out.

The second approach is based again on the Normalized Mutual Information. We used the same approach to compare clusters on the gene side and on the sample side. Given a partition in clusters of genes or samples the *NMI* was estimated as

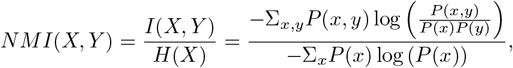

being *X* the set of cluster annotations (i.e. if *X* = {*c*_1_, *c*_1_, *c*_1_, *c*_2_ … *c*_*k*_} it means that the first three samples belong to cluster *c*_1_…). *P* (*x, y*) is the probability that a sample is classified in cluster *x* by the first algorithm and in cluster *y* by the second algorithm. *P* (*x*) is the probability that a sample belongs to cluster *x*, namely the size of cluster *x* divided by the number of samples. *NMI* reaches one if the two partitions are identical: one gains no information reading the second sequence when the first is known. Notice that this measure is not affected by a reshuffling of cluster labels which are completely arbitrary.

### 3.5 Gene selections

We used three different gene selection protocols.

#### Highly variable genes

this selection was performed using *scanpy* python package [60]. The package performs the analysis using the dispersion (variance over mean) [46]. Note that this kind of selection is always performed on the logarithmic data. See Figure S4 in Supplementary for an example of this kind of selection.

#### Housekeeping genes

These are typically constitutive genes that are required for the maintenance of basic cellular function. They are expressed in all cells of an organism under normal and pathophysiological conditions. We downloaded the list of 6 289 housekeeping genes provided by [9]. Among these, we filtered a total of 3 000 genes highly variable genes. The highly variable filter was performed taking into account only the housekeeping genes.

#### Random genes

These were selected using the `sample(random_state=42)` function provided by the *DataFrame* module of *pandas* package.

### 3.6 Topic distributions and fuzzy memberships

We are interested in studying the relevance of different topics in a given sample (or in a group of samples). To this end, we introduced a centered version of the *P* (topic|sample) distribution. Let us first recall the definition of *P* (topic|sample) in hSBM and LDA. In hSBM it is estimated as

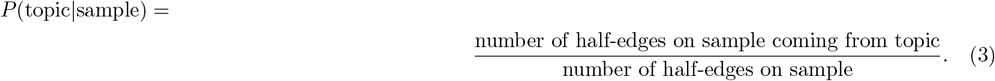

In LDA it is a *K* dimensional vector *θ*_*i*=1…*R*_ over topics (where *K* is the number of topics and *R* the number of samples), whose prior is a Dirichlet distribution *θ*_*i*_ = *Dir*(*α*) [65].

Topic models also allow to estimate the *P* (gene|topic) distributions that quantify the contribution of each gene to a topic. In hSBM, this is estimated as:

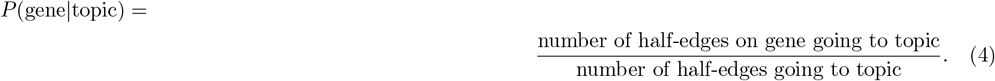

In LDA it is a *N* (number of genes) dimensional vector *ϕ*_*k*=1…*K*_ over genes, whose prior is a Dirichlet distribution *ϕ*_*k*_ = *Dir*(*β*).

### 3.7 Box topic and gene ontologies

Starting from *P* (topic|sample) we constructed a centered versions defined as:

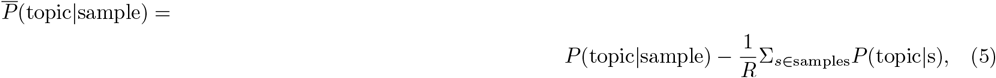

being *R* the total number of samples. This centered 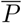 (topic|sample) can be represented with a box plot, after grouping samples by their tissue. These are the box plots reported in Fig. 7

Topics are nothing but lists of genes, which are naturally brought to perform gene ontology tests. We searched for enrichment using GSEA [53].

### 3.8 Cancer and healthy samples in a unified dataset

In the latest analyses, we processed data of samples unified from GTEx and The Cancer Genome Atlas (TCGA), this dataset was prepared and described in [57]. Normalized data were downloaded from [58].

We filtered samples in the analyses described in this manuscript, in particular, we sorted tissues by the number of samples available and got the 8 tissues with the largest number of samples: Breast, Colon, Esophagus, Liver, Lung, Prostate, Stomach, and Uterus. We randomly selected 100 samples per tissue which we used to define the topic space, while all the other samples were used for training and validation. We set a cutoff above 100 000 FPKM to reduce the number of edges and avoid overflows during the fitting procedure. This clipped the 0.2% of the links.

We collected the metadata through the GTEx portal and using the Genomic Data Commons tools [14] for the TCGA samples.

### 3.9 Predictor on latent space

We built a Neural Network predictor using topics as features, in other words, we trained a simpler model on a low-dimensional space instead of a complex model on the original ∼20 000 dimensional space.

The first step of this analysis was picking up 800 samples of 8 tissues from the unified dataset [58]. We fit the hierarchical Stochastic Block Model using this subset. In output, we obtained the topic distribution of samples.

At this point, we considered topics as features and *P* (topic|sample) as entries of the design matrix *X*. The entry *X*_*ij*_ is the *P* (topic_*j*_ |sample_*i*_). The matrix needed further normalization to be fitted using Stochastic Gradient Descent later. The normalized matrix 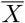 was obtained by subtracting features means and dividing each feature by its range. The entries of this new matrix are 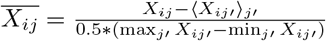.

The dataset was split into a training and a test set; the training set contains the 95% of the samples. This is quite unbalanced, but we were not going to use it to evaluate the performance of the model. Moreover, 25% of the training set was used as the validation set. The model consisted of a neural net with one hidden layer with 100 neurons activated by a *ReLU* function, the optimizer was set to Stochastic Gradient Descent (SGD). Finally, the output layer used a *softmax* activation to classify tissues.

We wanted to evaluate the performance on unseen samples, never retraining either topic modeling or the neural net. From the original dataset, we selected all the samples not involved in topic modeling. These were ∼5 000 new samples never fitted by hSBM neither by the neural net. We projected this *unseen* samples into the topic space. To do this, we firstly selected the genes involved in topic modeling, the ones that passed the highly variable filter we imposed. The *P* (topic|sample) for the new samples were simply *P* (topic|sample) = Σ_gene_*P* (topic|gene) \**P* (gene|sample), in other words each expression array is multiplied to the matrix *genes*×*topics*. We now evaluate the Neural Network performances in separating tissues on this new dataset and obtained an accuracy of 0.9273. The Area Under Curve score is here 0.9852.

A similar experiment can be done, but considering “healthy” and “diseased” as labels. In this case, the output layer is simply a single neuron activated by a *sigmoid* function. The scores when predicting the status of the samples are 0.9474 for accuracy and 0.9762 for AUC.

The confusion matrix and the Receiving Operating Characteristic curves (reported in Figure 10) were estimated using scikit learn [32]. The ROC curve represents the True Positive Rate or sensitivity 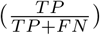 versus the False Positive Rate or 1 –specificity 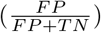.

The whole predictor model was implemented using keras [7] and using TensorFlow as the backend.

We measured the K-NN performance using the accuracy and the AUC scores. We applied K-NN setting `n_neighbors` to 5 and using *euclidean* `metric`.

### 3.10 Code availability

Code and notebooks to reproduce our analyses are available from https://github.com/BioPhys-Turin/ topics, Topic and Cluster names (i.e. Cluster 1, Topic 1) may vary between the draft and the repository. We changed them for writing purposes.

## 4 Authors contribution

conceptualization, F.V., M.O. and M.C.; methodology, F.V., M.O. and M.C.; software, F.V.; visualization, F.V.; writing–preparation original draft, F.V. and M.O.; writing–review and editing, F.V., M.O. and M.C.;

## 5 Acknowledgements

The results shown here are in part based upon data generated by the TCGA Research Network: https://www.cancer.gov/tcga.

We want to acknowledge the Competence Centre for Scientific Computing C^3^S which provided us access to the computing cluster OCCAM.

## Footnotes

1 the NMI* top scores (hSBM 43, LDA 47, TM 46, WGCNA 27 and hierarchical 35) are slightly smaller than the scores obtained with the standard gene selection presented in Figure 1).

2 This analysis cannot be done with hierarchical clustering since the original points are not projected in a low-dimension space.

