## Supplementary Material for "Exploring the latent space of transcriptomic data with topic modeling"

### Exploring the gene expression latent space: a topic modeling approach

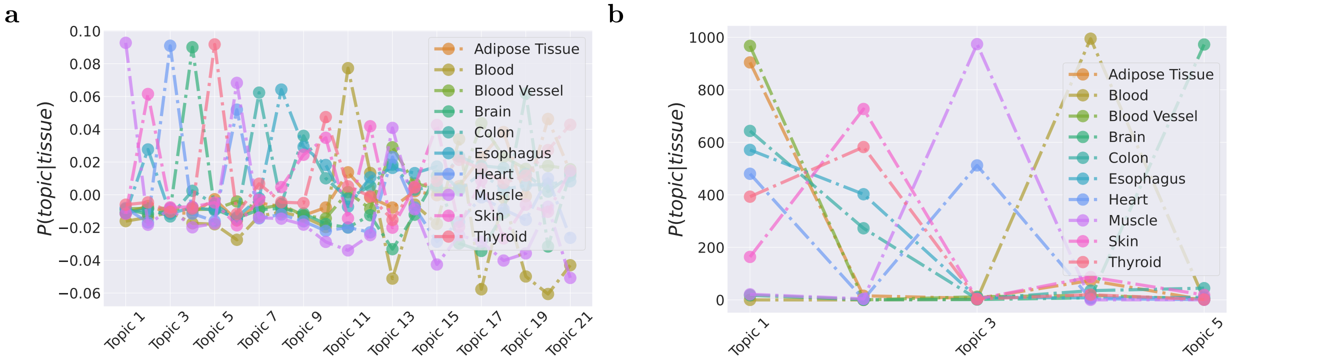


Figure S1: We reported $P(\text{topic}|\text{tissue})$ ($P(\text{topic}|\text{sample})$ averaged over tissues) for different tissues using the projections provided by (a) WGCNA and (b) Topic Mapping.

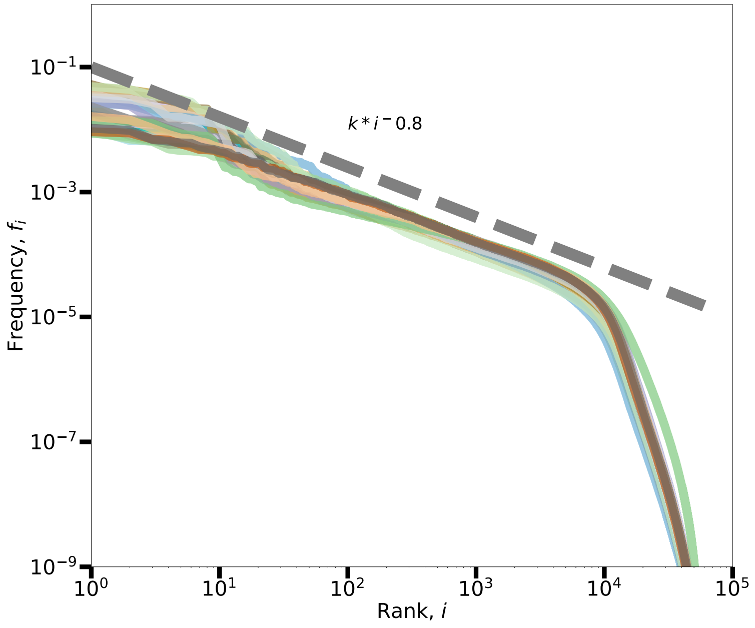


Figure S2: Rank plot of genes abundances in GTEx. The frequency $f_i$ is estimated as $f_i = \frac{\Sigma_{s=1...R} n_{gs}}{\Sigma_{s=1...R} M_s}$ being $M_s = \Sigma_{g=1...N} n_{gs}$, $R$ the number of samples and $N$ the number of genes.

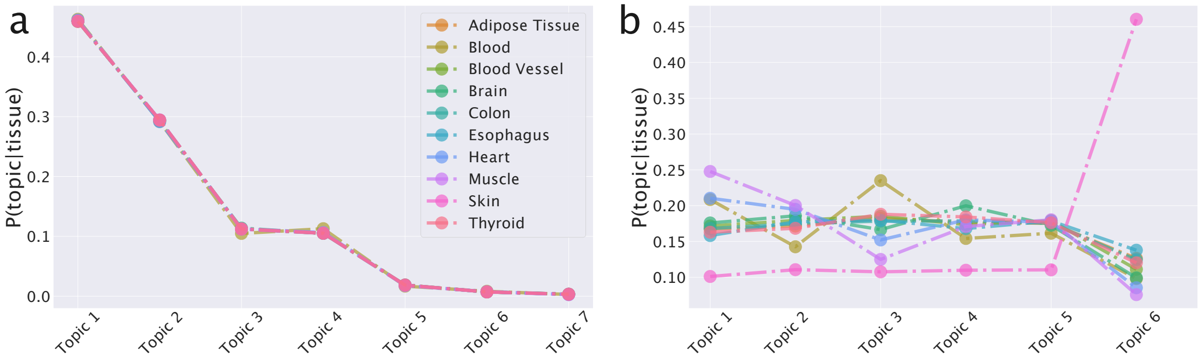


Figure S3: We reported $P(\text{topic}|\text{tissue})$ for different tissues for (a) hSBM and (b) LDA in experiments considering only housekeeping genes.

| algorithms | | NMI |
|---|---|---|
| hsbm | tm | 0.256 |
| hsbm | lda | 0.070 |
| hsbm | wgcna | 0.058 |
| tm | lda | 0.150 |
| tm | wgcna | 0.656 |
| lda | wgcna | 0.147 |

Table S1: *NMI* between the genes mixtures in different algorithms (hierarchical doesn't provide information on genes).

| algorithms | | NMI |
|---|---|---|
| hsbm | tm | 0.573 |
| hsbm | lda | 0.806 |
| hsbm | wgcna | 0.772 |
| hsbm | hierarchical | 0.867 |
| tm | lda | 0.659 |
| tm | wgcna | 0.707 |
| tm | hierarchical | 0.610 |
| lda | wgcna | 0.836 |
| lda | hierarchical | 0.852 |
| wgcna | hierarchical | 0.813 |

Table S2: *NMI* between the clusters mixtures in different algorithms.

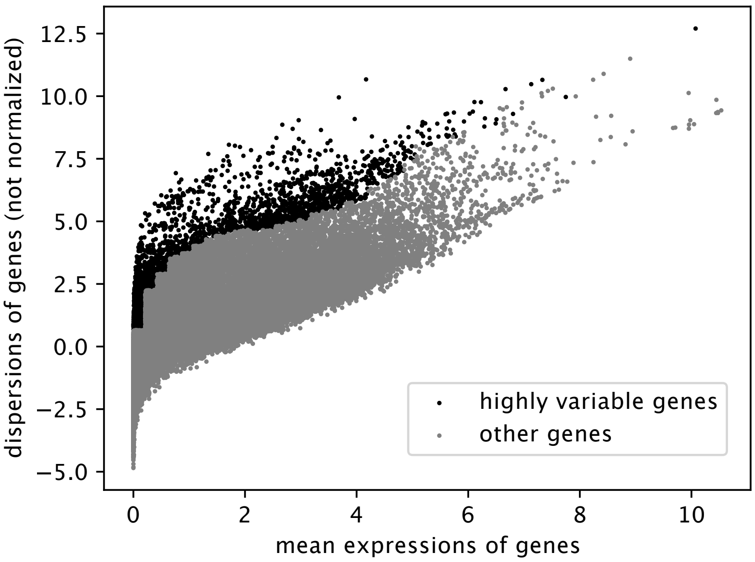


Figure S4: We use *scanpy* standard tool to select the highly variable genes. The dispersion is defined as $\frac{\sigma^2}{\text{mean}}$ and it scales with mean. Highly variable genes are the most dispersed in each bin of the mean. We used `scanpy.pp.highly_variable_genes(adata, n_top_genes = 3000, n_bins = 50)`

#### Latent spaces

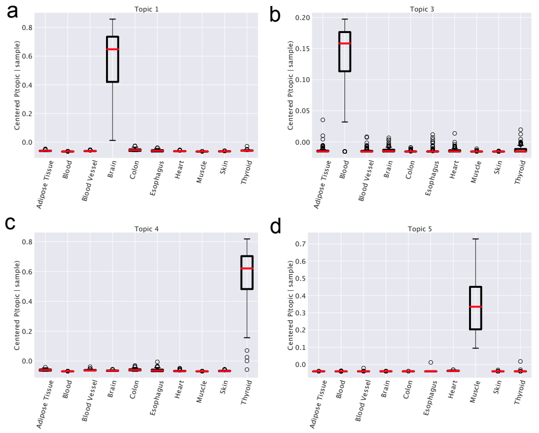


| Term | FDR q-value |
|---|---|
| Topic 1 (20) | |
| GO_ANTIMICROBIAL_HUMORAL_RESPONSE | $6.44e-5$ |
| CHARAFE_BREAST_CANCER _BASAL_VS_MESENCHNCHYMAL_UP | $6.44e-5$ |
| Topic 3 (20) | |
| GSE22886_NAIVE_BCELL_VS_NEUTROPHIL_DN | $1.27e-14$ |
| GSE29618_MONOCYTE_VS _PDC_DAY7_FLU_VACCACCINE_UP | $1.41e-12$ |
| Topic 4 (20) | |
| GO_EPIDERMIS_DEVELOPMENT | $7.89e-23$ |
| GO_SKIN_DEVELOPMENT | $5.85e-17$ |
| Topic 5 (20) | |
| GSE22886_NAIVE_BCELL_VS_NEUTROPHIL_DN | $9.66e-12$ |
| MODULE_84 | $4.11e-8$ |

Figure S5: Gene ontologies enrichment test performed on different LDA's topics using GSEA. In brackets the number of genes.

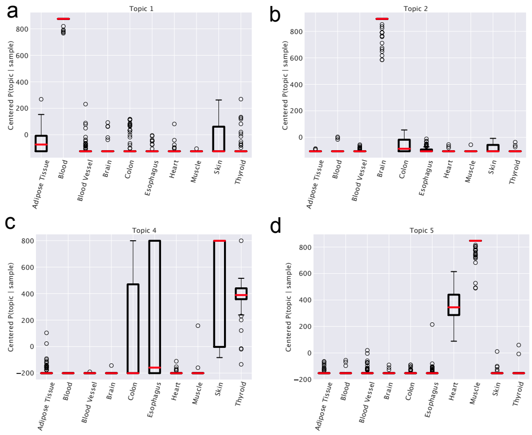


| Term | FDR q-value |
|---|---|
| Topic 1 (26) | |
| CHEN_METABOLIC_SYNDROM_NETWORK | $1.92e-7$ |
| RODWELL_AGING_KIDNEY_UP | $3.3e-7$ |
| Topic 2 (24) | |
| MODULE_12 | $1.79e-3$ |
| GSE45365_NK_CELL_VS_CD8A_DC_DN | $1.79e-3$ |
| Topic 4 (25) | |
| HOLLERN_EMT_BREAST_TUMOR_DN | $2.51e-22$ |
| ONDER_CDH1_TARGETS_2_DN | $6.85e-19$ |
| Topic 5 (23) | |
| GO_CONTRACTILE_FIBER | $1.4e-26$ |
| GNF2_MYL2 | $5.85e-19$ |

Figure S6: Gene ontologies enrichment test performed on different TM's topics using GSEA. In brackets the number of genes. TM outputs 5 topics, they are not enough to assign a topic to each tissue.

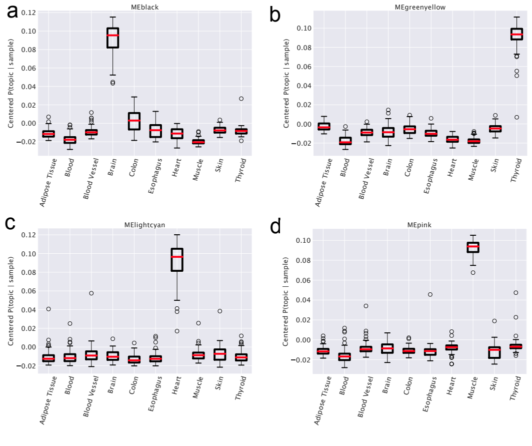


| Term | FDR q-value |
|---|---|
| Topic black (389) | |
| GO_SYNAPSE | $4.64e-78$ |
| GO_NEURON_PROJECTION | $1.19e-49$ |
| Topic greenyellow (328) | |
| RODRIGUES_THYROID_CARCINOMA _POORLY_DIFDIFFERENTIATED_DN | $7.68e-27$ |
| RODRIGUES_THYROID_CARCINOMA _ANAPLASTICTIC_DN | $1.2e-22$ |
| Topic lightcyan (56) | |
| GO_CONTRACTILE_FIBER | $4.18e-31$ |
| GO_MUSCLE_SYSTEM_PROCESS | $2.99e-26$ |
| Topic pink (192) | |
| GO_CONTRACTILE_FIBER | $2.71e-66$ |
| HALLMARK_MYOGENESIS | $8.52e-61$ |

Figure S7: Gene ontologies enrichment test performed on different WGCNA's topics using GSEA. In brackets the number of genes, in this setting the number of genes is hundreds, this can bias the hypergeometric P-value.

#### Scores

In the paragraph about classifying samples with neural networks, we tested three settings:

- samples used to train topic modeling;
- samples projected in the topic space (these points are not used to train topic modeling);
- samples directly from the dataset without any processing;

All the results for a Neural Network NN and for k-NN are reported in the following tables.

| data space | log-transformed data space | unseen data |
|---|---|---|
| | Primary site (tissue) | |
| acc: 0.8375 auc: 0.838 | acc: 0.969, auc: 0.983 | acc: 0.820 auc: 0.902 |
| | Status (healthy or diseased) | |
| acc: 0.8375 auc: 0.8377 | acc: 0.9875 auc: 0.9839 | acc: 0.9226 auc: 0.9268 |
| | All healthy primary sites | |
| acc: 0.9362 auc: 0.9659 | acc: 0.9804 auc: 0.9896 | n.a. |

Table S3: Scores using the K-NN model. The rows represent the three different datasets discussed in the main text: 10 healthy tissues, unified healthy and diseased tissues, and all GTEx tissues. The scores in the first two columns are estimated in the original data space before and after applying the log-transformation. The last column represents samples not used for training topic modeling.

| test set | unseen data |
|---|---|
| Primary site (tissue) | |
| acc: 1 auc: 1 | acc: 0.9273 auc: 0.9852 |
| Status (healthy or diseased) | |
| 0.9750 auc: 1 | acc: 0.9474 auc: 0.9762 |
| All healthy primary sites | |
| acc: 0.9333 auc: 0.9980 | n.a. |

Table S4: Neural Network scores in different sets. The rows represent the three different datasets: 10 tissues, Healthy and diseased tissues, and all GTEx tissues. The first column represents the test error when using samples passed through topic modeling. The second column represents the test error when using points just projected into the topic space not being used for training the topic model.

| | Homogeneous | Not homogeneous |
| --- | --- | --- |
| Complete | 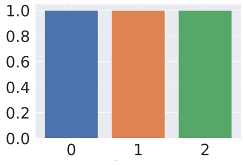 | 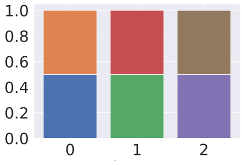 |
| Not complete | 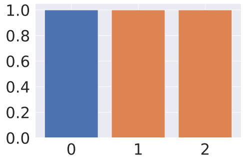 | 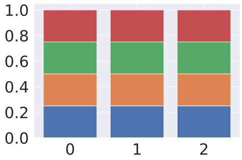 |

Table S5: Examples of homogeneity and completeness. Homogeneous clusters contain all nodes with the same label. A label is complete if it is fully represented by a single cluster. In this image some examples of these definitions. The *NMI* score discussed in this work is nothing but the geometric average of completeness and homogeneity.
